# Genomic prediction with allele dosage information in highly polyploid species

**DOI:** 10.1101/2021.06.22.449437

**Authors:** Lorena G. Batista, Victor H. Mello, Anete P. Souza, Gabriel R. A. Margarido

## Abstract

Several studies have shown how to leverage allele dosage information to improve the accuracy of genomic selection models in autotetraploids. In this study we expanded the methodology used for genomic selection in autotetraploids to higher (and mixed) ploidy levels. We adapted the models to build covariance matrices of both additive and digenic dominance effects that are subsequently used in genomic selection models. We applied these models using estimates of ploidy and allele dosage to sugarcane and sweet potato datasets and validated our results by also applying the models in simulated data. For the simulated datasets, including allele dosage information led up to 140% higher mean predictive abilities in comparison to using diploidized markers. Including dominance effects was highly advantageous when using diploidized markers, leading to mean predictive abilities which were up to 115% higher in comparison to only including additive effects. When the frequency of heterozygous genotypes in the population was low, such as in the sugarcane and sweet potato datasets, there was little advantage in including allele dosage information in the models. Overall, we show that including allele dosage can improve genomic selection in highly polyploid species under higher frequency of different heterozygous genotypic classes and high dominance degree levels.

## Introduction

Polyploids are organisms with more than two sets of chromosomes. The number of sets of chromosomes in an organism is named its ploidy level. Polyploids are classified into two major categories of auto and allopolyploids. Allopolyploids result from the combination of distinct parental genomes and are characterized by preferential pairing of chromosomes, with bivalent chromosome formation in meiosis and disomic inheritance at each locus. In contrast, autopolyploids have more than two homologs per homology group, often leading to the formation of multivalent chromosomes and polysomic inheritance (Soltis and Soltis, 2000).

Many economically important species are autopolyploids. Among these, a high ploidy level (>4) is observed in a number of species such as sweet potato, sugarcane, and some ornamental flowers and forage crops. Sweet potato, an autohexaploid, is the fourteenth most important food crop in the world regarding production volume (FAOSTAT, 2020), and sugarcane, with ploidy levels ranging up to 16 (Garcia *et al*. 2013), accounts for 80% of the worldwide sugar production (CIRAD) and has potential to become the main crop for bioenergy production. The main bottleneck in breeding programs for these species is the long process for selection of cultivars. A traditional sugarcane breeding program is usually divided in several phases of selection, each consisting of large experiments that are usually conducted for more than one crop cycle (Cheavegatti-Gianotto *et al*. 2011; Zhou 2013), taking up to 12 years from the initial crosses until commercial cultivar release (Park *et al*. 2007). Sweet potato breeding programs follow a similar breeding scheme, with selection of cultivars taking up to 10 years (Katayama *et al*. 2017). In this context, there is a pressing need for the deployment of strategies to reduce experimental costs and time for selection of cultivars.

Genomic selection is a viable way of achieving improvement in breeding programs in terms of time and costs (Heffner *et al*. 2009). Genomic selection consists of using a representative population that is both genotyped and phenotyped (i.e., the training population) to predict the effect of genetic markers widely spread throughout the genome. The predicted effects are then used to predict the breeding or genotypic value of genotyped individuals (Meuwissen *et al*. 2001). This allows selection to be carried based on predicted breeding values, reducing the need for further costly phenotypic evaluations and shortening the time needed for selection of the best genotypes. Genomic selection has been successfully implemented in several crop breeding programs (Bernardo and Yu 2007; Heffner *et al*. 2009; Crossa *et al*. 2010; Resende *et al*. 2012; Duhnen *et al*. 2017) and can potentially increase genetic gain in sugarcane breeding programs (Voss-Fels *et al*. 2021; Hayes *et al*. 2021). Although genomic selection can greatly improve breeding programs, its implementation demands a relatively large set of genetic markers to be consistently obtained at feasible costs, a process which is hindered in complex genomes such as those of highly autopolyploid species.

Due to the complexity of their genomes, genetic studies in autopolyploid species were historically mostly carried using either dominant or diploidized markers (Dufresne *et al*. 2014), that is, polymorphisms that are either detected in a presence/absence fashion or polymorphisms where all heterozygous genotypes are collapsed into a single class. When using only dominant or diploidized markers, information on the different categories of heterozygous genotypes is effectively lost. However, several new tools are now available that allow estimating the allele dosage (i.e., the quantitative genotypes) of markers (Serang *et al*. 2012; Blischak *et al*. 2018; Gerard *et al*. 2018; Clark *et al*. 2019), and information of all possible genotypic classes can now potentially be used in genomic studies of polyploids.

In autotetraploids, several studies have shown how to leverage allele dosage information to improve the accuracy of genomic selection models (Slater *et al*. 2016, 2016; de Bem Oliveira *et al*. 2018; Hawkins and Yu 2018; Endelman *et al*. 2018; Amadeu *et al*. 2020). However, to our knowledge no studies so far have expanded these methodologies to specifically address organisms with higher ploidy levels. In this paper, we generalize genomic selection models used in autotetraploids and assess the accuracy of genome-wide prediction when incorporating allele dosage information in sugarcane and sweet potato datasets, two highly autopolyploid species. In order to validate our results, we also assess the accuracy of prediction in four simulated datasets.

## Material and Methods

### 1. Genetic material and field experiments

The sugarcane dataset consisted of a segregating F_1_ progeny of 179 individuals derived from the crossing of two commercial cultivars, IACSP95-3018 (female) and IACSP93-3046 (male). The first field experiment was set in Sales de Oliveira, SP, Brazil, in 2007. A randomized complete block design with four replicates was used and evaluations were carried in the harvest years of 2008 (plant cane) and 2009 (ratoon cane). The full-sib progeny was then clonally propagated for the second field experiment that was set in Ribeirão Preto, SP, Brazil, in 2011. A randomized complete block design with three replicates was used and evaluations were carried in 2012 (plant cane), 2013 and 2014 (ratoon cane). Both parents were included in each block of the two experiments. All replicates were used to collect phenotypes for stalk diameter (cm), stalk height (cm) and stalk weight (kg) in both experiments. Also, two blocks in each experiment were used to collect phenotypes for soluble solids content (Brix), sucrose content and fiber percentage.

The sweet potato dataset consisted of phenotypic records on 282 accessions of *Ipomoea batatas* made available by Jackson *et al*. (2018), which are part of a broader group of 731 accessions randomly selected from the USDA germplasm bank in Griffin, Georgia, United States. These materials have origin in more than 30 countries in eight geographic regions (Africa, Australia, Caribean, Central America, East Asia, North America, Pacific islands and South America). The accessions were planted in field trials and phenotyped in the years 2012, 2013 and 2014. In in this study, we only used phenotypic data from the stele colorimetry analysis. The stele colorimetry data included values of the green-red coordinate (**a**), the yellow-blue coordinate (**b**), colour saturation (**C**), lightness (**L**), and hue angle (**h**).

### 2. Genotyping

For the sugarcane population, parents and F_1_ progeny were genotyped using the genotyping-by-sequencing protocol of Elshire *et al*. (2011). Reduced representation libraries were prepared using the PstI restriction enzyme. PstI is a rare-cutting enzyme, because its restriction site has a length of 6 bp, allowing a higher genotyping depth (Poland and Rife 2012). Four lanes containing 96-plex libraries were sequenced using the Illumina GAIIx and, subsequently, another four lanes with the same 96-plex libraries were sequenced using the Illumina NextSeq500 platform.

The genotyping-by-sequencing protocol used for the sweet potato accessions is described by Wadl *et al*. (2018), where a modified genotyping-by-sequencing protocol optimized for highly heterozygous and polyploid genomes was used (GBSpoly). They used a combination of *Cvi*AII and *Tse*I restriction enzymes for preparing the libraries (restriction sites with 4 and 5bp, respectively). Libraries were multiplexed with 96 pooled samples. In this study, we used the raw read data the authors in Wadl *et al*. (2018) made available in the NCBI database with accession code SRP152827.

For the sugarcane dataset, we called variants using a modified version of the TASSEL-GBS pipeline (Pereira *et al*. 2018). This version provides exact read counts of the alleles at each SNP locus. We used default values in all plugins of the pipeline, except for the MergeDuplicateSNPs plugin, in which we used the argument *callHets* and set the *misMat* argument value to 0.3. These values were chosen to allow a greater number of heterozygous SNP loci to be kept in subsequent steps. The sequenced reads were then aligned to the methyl-filtrated assembly of the sugarcane genome (Grativol *et al*. 2014), using the software Bowtie2 (Langmead and Salzberg 2012).

The sweet potato raw reads were first aligned to the two ancestral reference genomes *I. trifida* and *I. triloba* (Shiotani 1988; Oracion *et al*. 1990; Freyre *et al*. 1991) using Bowtie2 (Langmead and Salzberg 2012). We then used the HaplotypeCaller tool in the GATK software (version 4.1.4) to call SNPs, indels and copy number variants.

For both species we used the read count information of each SNP to estimate their ploidy level and call sample genotypes using the software SuperMASSA and VCF2SM (Serang *et al*. 2012; Pereira *et al*. 2018). For sugarcane, ploidy levels ranging from two to 20 were evaluated and only SNPs with ploidy estimates between six and 14 were kept (Garcia *et al*. 2013). We also filtered for a minimum mean read depth per individual of 50 reads, maximum mean read depth per individual of 500 reads, minimum posterior probability of genotype configuration of 0.8, minimum posterior probability of each genotype assignment of 0.5, and minimum call rate of 50%. For sweet potato, ploidy levels ranging from four to eight were evaluated and only SNPs with a ploidy estimate of six were chosen. We used a minimum mean read depth per individual of 45 reads, maximum mean read depth per individual of 200 reads and the remaining arguments were the same as for sugarcane.

We used the R package *updog* (Gerard *et al*. 2018) to reestimate the genotypes of the SNPs that met the filtering criteria in both species. The *updog* package has the advantage of accounting for allelic bias, overdispersion and sequencing errors when estimating SNP genotypes, given a predetermined ploidy level. For sweet potato, SNP sets resulting from the alignment with each of the reference genomes were merged, and redundant SNPs (i.e., with identical genotype calls for all individuals) were removed.

Finally, we performed a chi-squared segregation test on the population genotype class frequencies. For the sugarcane F_1_ progeny, based on the estimates of SNP genotypes in the parents, we tested the goodness-of-fit of marker genotypes to a hypergeometric distribution of gametes (Mollinari and Serang 2015). For the sweet-potato diversity panel we tested the goodness-of-fit of marker genotypes to the distribution expected under Hardy-Weinberg equilibrium. Using the Bonferroni correction, only SNPs with p-values greater than a 5% threshold were kept.

### 3. Phenotypic mixed model analysis

Adjusted phenotypic means (i.e., BLUEs - best linear unbiased estimates) for each individual were obtained using a two-stage analysis (Damesa *et al*. 2017). All analyses were performed using ASReml-R (Butler *et al*. 2009). Stage one consisted of a within-site analysis, where the genotype effect was considered fixed and the remaining effects were considered as random (harvest effects, blocks-within-harvest effects, and genotype × harvest interaction effects). The covariance matrix (Ω_***j***_) for the vector of genotype effects (***û***_***j***_) in site *j* was obtained from the inverse of the coefficient matrix of the mixed model equations, returned as *Cfixed* in the asreml object (Endelman *et al*. 2018). Stage two was a joint analysis considering the two sites, using the following linear model:

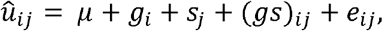

where *û*_*ij*_ is the genotype effect estimate obtained in the stage one analyses, the parameter *µ* is the intercept, *g*_*i*_ is a fixed effect of genotypes, *s*_*j*_ is a random effect of sites, (*gs*)_*ij*_ is a random effect for the genotype × site interaction, and the variance of the residual *e*_*ij*_ is (*ω* ^*ij*^)^-1^, where *ω*^*ij*^ is the *i*th diagonal element of Ω_*j*_ ^-1^ from the stage one analysis (Damesa *et al*. 2017). The BLUEs of the genotypes obtained after this stage were subsequently used to fit the genomic selection models.

### 4. Genomic selection models

We incorporated allele dosage information in our genomic selection models by expanding and adapting the GBLUP methodology for autotetraploid species proposed by Endelman *et al*. (2018). In sugarcane, besides the higher ploidy, the model also has to account for different ploidy levels among SNP loci. In order to achieve that, we expanded the theory by adapting the estimation of the genomic covariance matrix of both the additive values (**G**) and the digenic dominance values (**D**).

Genomic predictions were obtained using the following linear model:

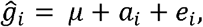

where *ĝ*_*i*_ is the BLUE of the *i*th individual obtained with the two-stage phenotypic analysis, *µ* is the intercept, a_*i*_ is the random effect of genotypes, and *e*_*i*_ is the random residual effect.

We used two covariance structures in the genomic selection model: i) **I**V_*r*_ + **G**V_*a*_, and ii) **I**V_*r*_ + **G**V_*a*_ + **D**V_*d*_, where I is the identity matrix, V_*r*_ is the residual variance, V_*a*_ is the additive genetic variance, and V_*d*_ is the dominance genetic variance. All analyses were performed using ASReml-R (Butler *et al*. 2009).

#### 4.1 Genomic covariance matrix of additive values (G)

Consider a matrix **X** with *n* rows and *m* columns, the rows corresponding to the individuals in the population and the columns corresponding to SNP loci, where each element *x*_*ij*_ corresponds to the dosage of the alternative allele for the *j*-th SNP in the *i*-th individual. If *p*_*j*_ is the frequency of the alternative allele at the *j*-th locus, we can obtain an *n* × *m* matrix **P** where the values in the *j*-th column all correspond to *p*_*j*_. For hexaploid sweet potato, subtracting 6**P** from **X** results in the matrix **W** of centered genotypes. The **G** matrix is then obtained by the formula:

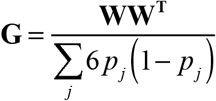

For sugarcane, because the SNPs have different ploidy levels, the same value of allele dosage for one SNP does not represent the same genotype for other SNPs with different ploidies. For example, for a hexaploid SNP an allele dosage value of six represents a homozygous genotype, while for an octoploid SNP the same value represents one of the possible heterozygotes.

To account for the different ploidy levels between SNPs, we used the following formula:

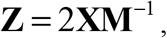

where **M** is an *m* × *m* diagonal matrix of ploidy values, such that each diagonal element *m*_*j*_ corresponds to the ploidy of the *j*-th SNP locus. The resulting matrix **Z**, with the same dimensions of **X**, has all its elements varying from 0 to 2, where 0 represents loci that are homozygous for the reference allele and 2 represents loci that are homozygous for the alternative allele, the values in between corresponding to heterozygous loci.

The subsequent steps to obtain **G** are the same as for diploids (VanRaden 2008). Subtracting 2**P** from **Z** results in the matrix **W** of centered genotypes. The **G** matrix is then obtained by the formula:

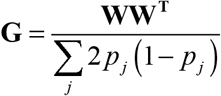

#### 4.2 Genomic covariance matrix of digenic dominance values (D)

We first introduce the expansion of the digenic dominance values in the autotetraploid model to a hexaploid scenario. Higher ploidy levels can be parametrized in a similar fashion. Considering a hexaploid SNP locus with two alleles B and b, the digenic effect for each allele pair can be obtained as demonstrated by Endelman *et al*. (2018), with the following set of equations:

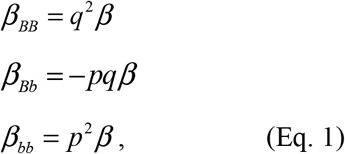

where *p* is the allele frequency of B, *q* is the allele frequency of b, with *q* = 1 − *p*, and *β* is the digenic dominance effect. Also, we have that:

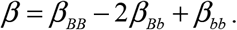

For a hexaploid locus, seven genotypic classes are possible in a population (i.e., allele dosages ranging from 0 to 6). For each genotypic class, different combinations of digenic effects are present. For example, for the genotypic class BBBBbb, there are 6 possible combinations of two B alleles, 8 possible combinations of a B allele with a b allele, and 1 possible combination of two b alleles. By replacing each digenic effect with their corresponding values in (Eq. 1), we obtain the total digenic dominance coefficient for each genotype class. Table 1 shows the combinations of digenic effects and the total digenic dominance coefficient for each possible genotype class of a hexaploid locus.

**Table 1.**
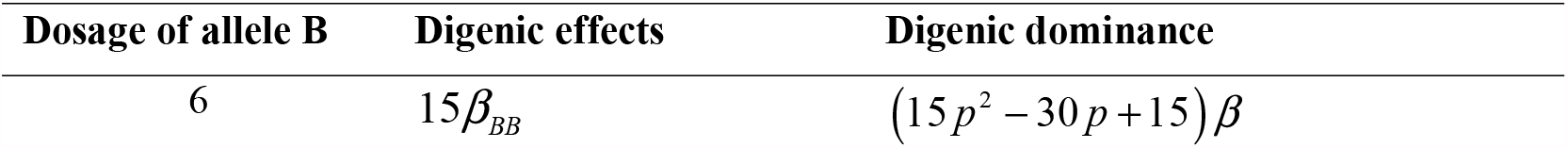
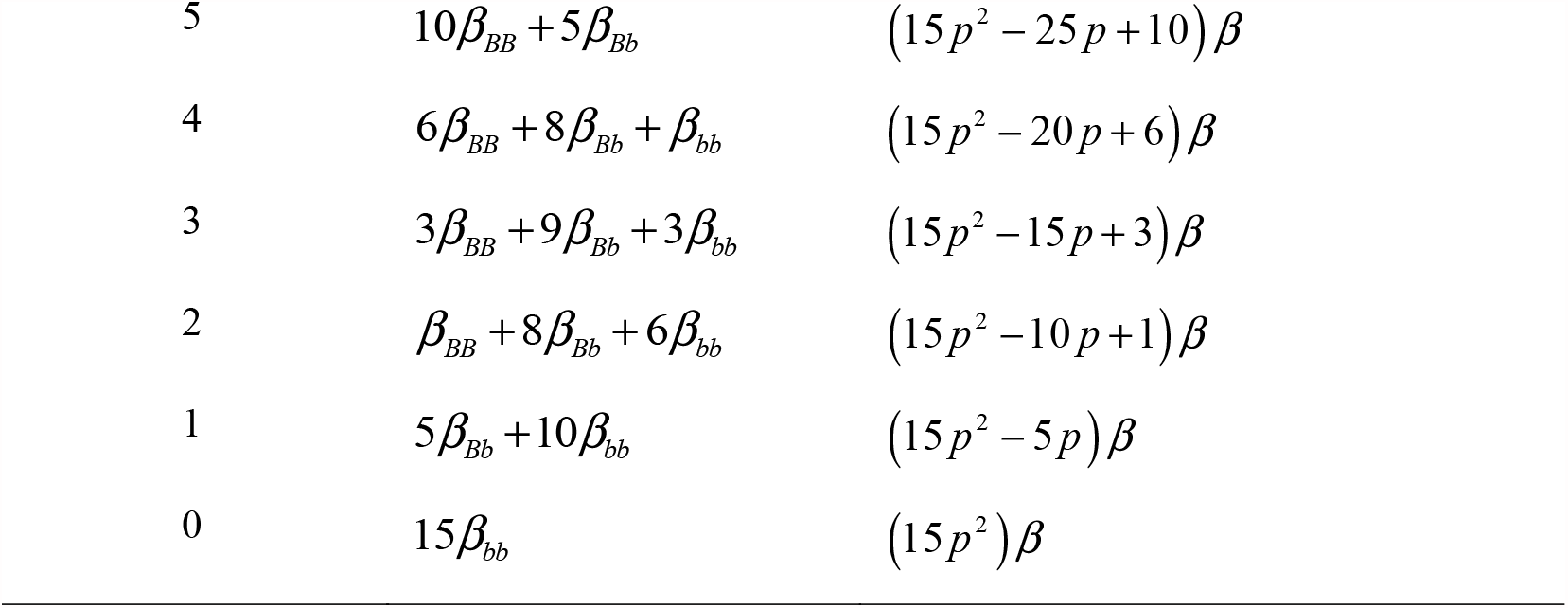
Digenic effects and total digenic dominance for each allele dosage level of a hexaploid locus with alleles B and b.

| Dosage of allele B | Digenic effects | Digenic dominance |
| --- | --- | --- |
| 6 | $15\beta_{BB}$ | $(15p^2 - 30p + 15)\beta$ |
| 5 | $10\beta_{BB} + 5\beta_{Bb}$ | $(15p^2 - 25p + 10)\beta$ |
| 4 | $6\beta_{BB} + 8\beta_{Bb} + \beta_{bb}$ | $(15p^2 - 20p + 6)\beta$ |
| 3 | $3\beta_{BB} + 9\beta_{Bb} + 3\beta_{bb}$ | $(15p^2 - 15p + 3)\beta$ |
| 2 | $\beta_{BB} + 8\beta_{Bb} + 6\beta_{bb}$ | $(15p^2 - 10p + 1)\beta$ |
| 1 | $5\beta_{Bb} + 10\beta_{bb}$ | $(15p^2 - 5p)\beta$ |
| 0 | $15\beta_{bb}$ | $(15p^2)\beta$ |

The formula to obtain the total digenic dominance for a given biallelic hexaploid locus can then be generalized as:

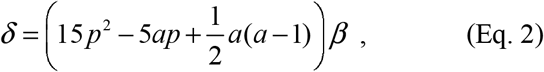

where *δ* is the total digenic dominance and *a* is the dosage of the B allele.

We used the same process described for hexaploid loci to obtain equations for other levels of ploidy. Table 2 shows the generalized formulas to obtain the total digenic dominance for even ploidies from six through 14.

**Table 2.** Formulas for the total digenic dominance for different levels of ploidy

| Ploidy | Total digenic dominance |
| --- | --- |
| 6 | $\left( 15p^2 - 5ap + \frac{1}{2}a(a-1) \right) \beta$ |
| 8 | $\left( 28p^2 - 7ap + \frac{1}{2}a(a-1) \right) \beta$ |
| 10 | $\left( 45p^2 - 9ap + \frac{1}{2}a(a-1) \right) \beta$ |
| 12 | $\left( 66p^2 - 11ap + \frac{1}{2}a(a-1) \right) \beta$ |
| 14 | $\left( 91p^2 - 13ap + \frac{1}{2}a(a-1) \right) \beta$ |

The formulas in Table 2 can then be generalized as:

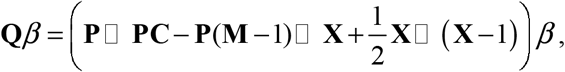

where □ represents the Hadamard product, **C** is an *m × m* diagonal matrix where each diagonal element *C*_*j*_ corresponds to 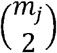, and **P, M** and **X** are as previously defined.

Finally, the genomic covariance matrix of digenic dominance values (**D**) was obtained with:

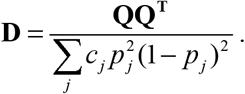

#### 4.3 Model and marker set comparisons

We compared two models for the genotype effects, one using only the additive **G** matrix (G model) and one using both the **G** and **D** matrices (G+D model). We also investigated the effect of using two different sets of genotypic information: i) a fully informative model considering SNP markers with ploidy and allele dosage estimates, and ii) diploidized SNP markers. The diploidized SNP set was obtained by setting the values of all heterozygous loci in matrix **Z** to 1. By doing so, all heterozygous genotypes were effectively merged in a single class, regardless of their dosage. For diploidized markers, the **G** and **D** matrices were obtained according to the established methodology commonly used for diploids (VanRaden 2008; Vitezica *et al*. 2013).

The models were compared in terms of predictive ability. For that, 1,000 cross-validation runs were carried out, such that in each run 10% of the population was sampled and used as the validation set, while the remaining 90% were used as the training set. We measured predictive ability as the correlation between predicted genotypic values and BLUEs of the individuals in the validation set.

### 5. Simulated datasets

#### 5.1 Population structure and founder genotypes

Stochastic simulations of two population structures were used to validate the accuracy of prediction of genomic selection models using allele dosage estimates for additive and dominance effects. One population was simulated with a nearly uniform distribution of all possible genotypic classes (Population 1). The second population was simulated with a higher frequency of simplex and homozygous genotypes which, in consequence, results in a higher prevalence of rare alleles (Population 2).

Genome simulation parameters were chosen to match the sweet potato genome. An autohexaploid genome consisting of 90 chromosomes (15 homology groups) was simulated and these chromosomes were assigned a genetic length of 1.43 Morgans and a physical length of 2×10^7^ base pairs (Wu *et al*. 2018). Sequences for each chromosome were generated using the Markovian Coalescent Simulator (Chen *et al*. 2009) and AlphaSimR (Gaynor *et al*. 2021). Recombination rate was inferred from genome size (i.e. 1.43 Morgans / 2×10^7^ base pairs = 7.15×10^−8^ per base pair), and the mutation rate was set to 2×10^−9^ and 2×10^−7^ per base pair for Populations 1 and 2 respectively. The probability of quadrivalent formation was set to 0.15 (Mollinari *et al*. 2020).

Simulated genome sequences were used to produce 50 founder genotypes. This was accomplished by randomly sampling gametes from the simulated genome to assign as sequences for the founders. Sites that were segregating in the founders’ sequences were randomly selected to serve either as causal loci or markers. For Population 1 we simulated a total of 1,000 segregating sites per homology group, of which 250 were selected as causal loci and 750 were selected as markers (3,750 causal loci and 11,250 markers in total). For Population 2 we simulated a total of 5,000 segregating sites per homology group, of which 250 were selected as causal loci and 750 sites with a high frequency of simplex and homozygous genotypes in the population were selected as markers. The allele frequencies and genotype distribution of markers in both populations are shown in Fig S1.1 and Fig S1.2 of Supplementary Material 1.

#### 5.2 Phenotype simulation

AlphaSimR defines an individual’s raw genotype dosage (*x*) as the number of copies of the “1” allele at a locus, which is then scaled in accordance with the ploidy level. The scaled dosages make inputs in the package invariant to ploidy level. The scaled additive genotype dosages (*x*_*A*_) are given by the formula:

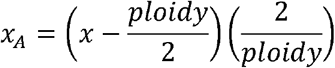

And the scaled dominance genotype dosages (*x*_*D*_) are given by the formula:

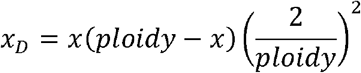

For autopolyploid organisms, this scaled dominance genotype dosage is consistent with the digenic dominance parametrization of the dominance model.

The true additive value of the simulated trait is then determined by the summing of its causal loci additive allele effects multiplied by the scaled additive genotype dosages. Additive allele effects were sampled from a standard normal distribution.

In the same way, the true dominance value of the simulated trait is determined by the summing of its causal loci dominance allele effects multiplied by the scaled dominance genotype dosages. The dominance effect (*d*) at a locus is the dominance degree (*δ*) at that locus times the absolute value of its additive allele effect (a):

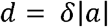

In this study, the dominance degrees were sampled from a normal distribution with variance 0.2 (Werner *et al*. 2020) and a mean of either 0.3 (low dominance) or 1 (high dominance). The additive and dominance effects were then scaled to achieve a desired genotypic variance of 1.

A phenotype was then simulated by summing the additive and dominance values and subsequently adding random error in order to achieve a heritability of 0.5.

#### 5.3 Population simulation

For each population structure (Populations 1 and 2) and dominance level (low dominance and high dominance) we simulated F_1_ populations with 300 individuals formed by randomly crossing the founder genotypes. Each of the four simulation scenarios (two populations x two dominance degree levels) was replicated 20 times. For each replicate, we deployed genomic selection models using a k-fold cross-validation scheme with k = 10. We measured predictive ability as the correlation between true and estimated genotypic values in the validation set.

## Results

We were able to obtain a large number of SNPs with estimates of ploidy and allele dosage in both sugarcane and sweet potato. In both species most of the genotypes were either homozygous or had only one copy of the alternative allele. The genomic selection models showed low prediction ability in the sugarcane dataset and moderate to high predictive ability in the sweet potato dataset. Overall the prediction ability values in both datasets showed little sensitivity to including ploidy and allele dosage information or dominance effects in the model. These results were replicated in simulated datasets where the marker genotype distribution was similar to the real datasets. In other simulated populations, which had a higher frequency of heterozygous markers, the highest values of predictive ability were achieved when including ploidy and allele dosage information in the models (full ploidy models). In these populations, including digenic dominance effects in full ploidy models was only advantageous when the dominance level was high. When using diploidized markers, including dominance effects increased predictive ability regardless of the dominance level.

### Genotyping

In sugarcane a total of 6,589 SNPs were kept after filtering for mean read depth, posterior probability of genotypes and ploidy estimates, call rate, and segregation distortion in the progeny. A total of 11 individuals did not have any sequenced reads and were considered not genotyped, thus being used in phenotypic analyses but not for genomic selection. A summary of ploidy and allele dosage estimates of the SNPs is shown in Fig. 1. The majority of the SNPs had ploidy estimates of ten (31.18%) and eight (28.93%), followed by 17.88% of SNPs with ploidy estimates of 12, 15.59% with an estimated ploidy of six, and 6.43% with ploidy 14. Within each ploidy level, most of the genotypes were either homozygous for the reference allele or had only one copy of the reference allele, with allele dosages of zero and one accounting for more than 50% of the total number of genotype calls for ploidy levels from six to 12. For ploidy 14, dosage estimates were more evenly distributed among different levels, but there was still an excess of dosages equal to zero and one.

**Fig 1.**
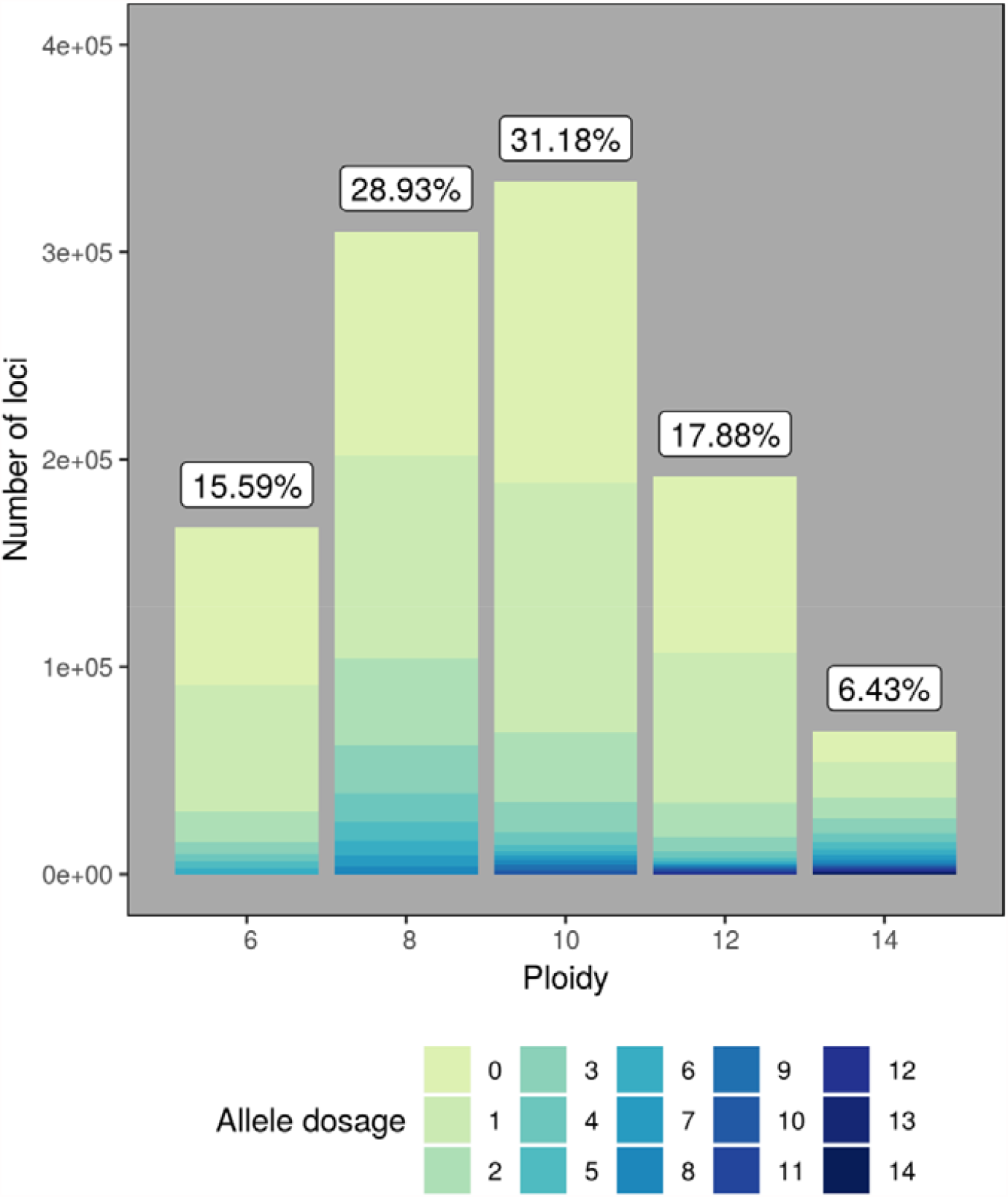
Summary of the estimates of ploidy and allele dosage for 170 sugarcane samples and 6,550 SNPs. The bars show the total number of loci per ploidy level, and different values of allele dosage are shown by different colours. For each ploidy level, the corresponding percentage of the total number of loci is shown above the bars.

In sweet potato we identified a total of 77,837 SNPs that were kept after filtering for mean read depth, posterior probability of genotypes and ploidy estimates, call rate, and segregation according to Hardy-Weinberg Equilibrium. A summary of allele dosage estimates of the SNPs is shown in Fig. 2. Most of the genotypes were either homozygous for the reference allele (53%) or had only one copy of the reference allele (13%), with allele dosages of zero and one (for both the reference and alternative alleles) accounting for more than 76% of the total number of genotype calls.

**Fig 2.**
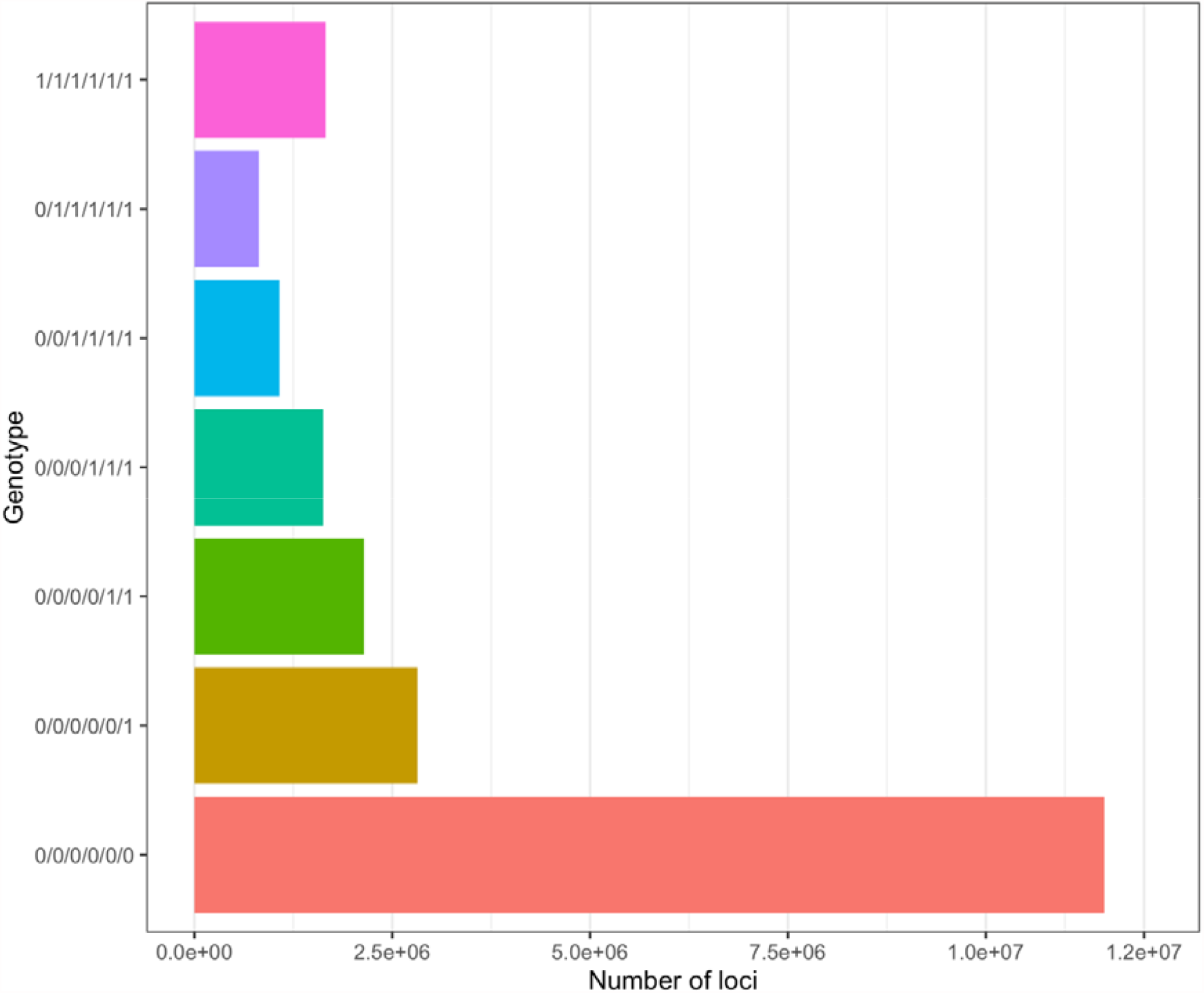
Genotype frequencies for 285 sweet potato samples and 77,837 SNPs. The bars show the total number of markers per genotypic class. Genotypic classes are shown with the alternative alleles represented as 1’s and the reference alleles represented as 0’s (e.g., “0/0/0/0/1/1” represents genotypes where the reference allele has a dosage of four and the alternative allele has a dosage of two).

### Genomic selection

#### Sugarcane

Overall, the predictive abilities of genomic selection in sugarcane were low, regardless of the model or marker set utilized. Fig. 3 shows the distribution of the predictive ability values in the sugarcane dataset over different cross-validation runs of the G and G+D models, when using all the makers with full ploidy and allele dosage information and using diploidized makers.

**Fig 3.**
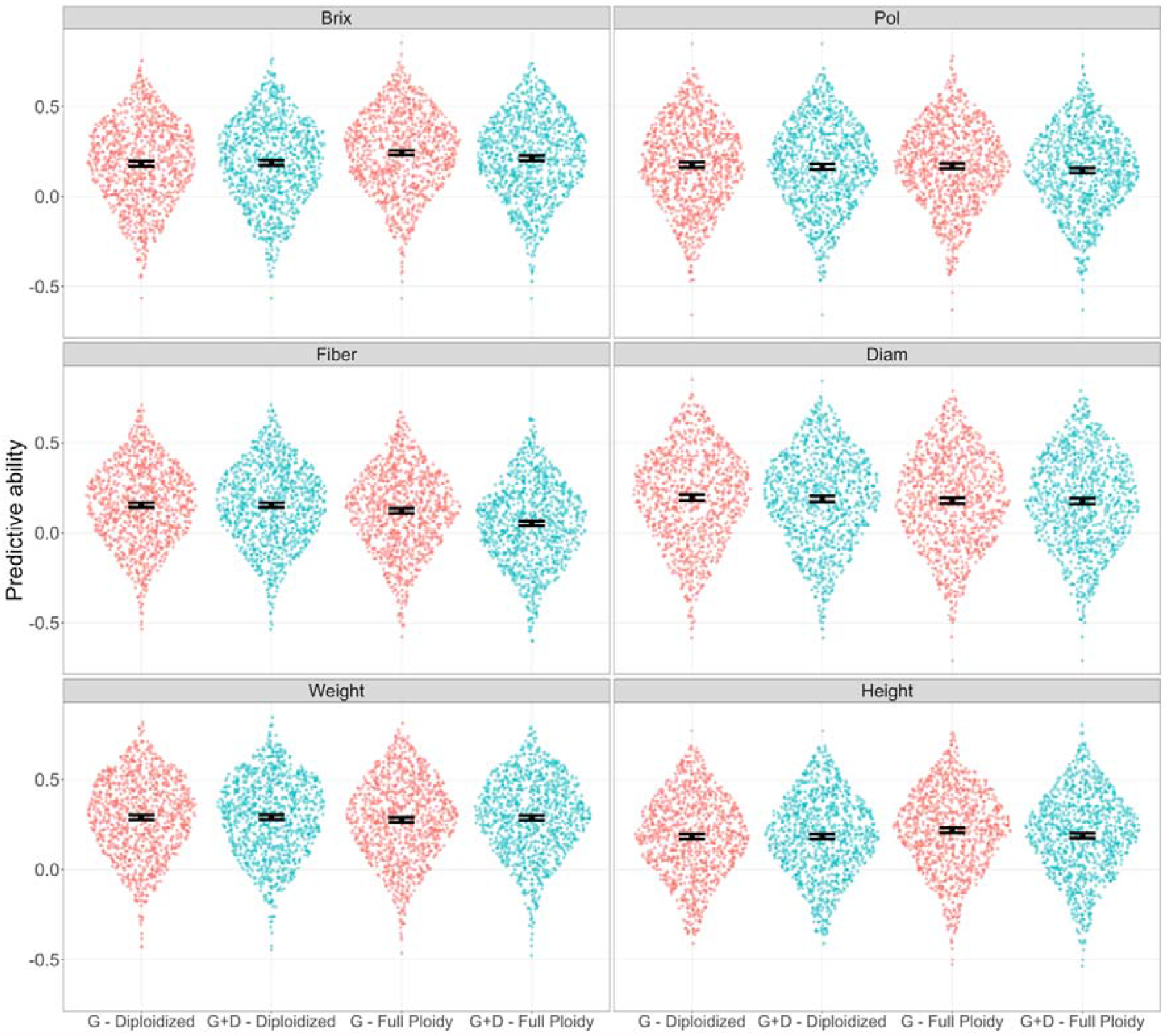
Distribution of the predictive ability values over different cross-validation runs of genomic selection in sugarcane. Values are shown when considering additive effects only (G) and considering additive and digenic dominance effects (G+D). Both models were compared when using markers with ploidy and allele dosage estimates (Full ploidy) and diploidized markers. The values are shown for traits soluble solids content (Brix), sucrose content (Pol), fiber percentage (Fiber), stalk diameter (Diam), stalk weight (Weight) and stalk height (Height). Mean and 95% confidence intervals are shown in black at the centre of each distribution.

For Brix, the G model using ploidy and allele dosage estimates showed the highest mean predictive ability (0.24), which was higher than that of the corresponding G+D model (0.21), and higher than the mean predictive abilities when using diploidized markers (0.18 for the G model and 0.19 for the G+D model). A similar pattern was observed for stalk height, where the G model using ploidy and allele dosage estimates had a mean predictive ability of 0.22, the full ploidy G+D model had a mean predictive ability of 0.19, and when using diploidized markers, the mean predictive ability did not exceed 0.18 for any of the two models.

For sucrose content, the G+D model had lower mean predictive abilities in comparison to the additive G model for all sets of markers, and the mean predictive abilities of the G model did not differ considerably between sets of markers. We observed a different pattern for stalk diameter, because the mean predictive ability of the G model when using ploidy and allele dosage estimates (0.18) was slightly lower than that achieved when using diploidized markers (0.20). With regard to the G+D model, the mean predictive abilities were equivalent for both sets of markers. A more marked difference between models was noticeable for fiber percentage, because for the full ploidy scenario the mean predictive ability of the G+D model (0.05) was much lower than for the G model (0.12). This, in turn, was lower than the mean predictive ability when using diploidized markers (0.15 for the G and G+D models). Lastly, for stalk weight, the mean predictive abilities were the highest among all traits, and the values did not differ significantly between models or sets of markers (ranging from 0.28 to 0.29).

In order to better understand the low values of predictive ability we observed in the sugarcane dataset, we performed a phenotypic variance partitioning analysis and obtained estimates of heritability for the evaluated traits (methodology details can be found in Supplementary Material 1). In general, the genotypic variance had a relatively small or intermediate magnitude for all the traits, with correspondingly low or intermediate heritability values. Fig. 4 shows the partitioning of the phenotypic variance into its main components. The residual variance had a large magnitude for all of the traits, corresponding to 36%, 35%, 49%, 58%, 48% and 34% of the phenotypic variation observed for Brix, sucrose content, fiber percentage, stalk diameter, stalk weight and stalk height, respectively. The effect of genotypes had an intermediate magnitude for stalk diameter and a small magnitude for the other traits, corresponding to 3%, 3%, 7%, 13%, 5% and 3% of the phenotypic variation observed for the same traits. The genotype × site interaction effect had an intermediate magnitude for fiber percentage, stalk diameter and stalk weight, with the variance due to the interaction component corresponding to, respectively, 13%, 15% and 10% of the observed phenotypic variation. For traits Brix, sucrose content and stalk height the variance due to the interaction component corresponded to 4%, 2% and 6% of the observed phenotypic variation, respectively. The heritability coefficients for traits Brix, sucrose content, fiber percentage, stalk diameter, stalk weight, and stalk height were 0.31, 0.35, 0.37, 0.55, 0.41 and 0.36, respectively.

**Fig 4.**
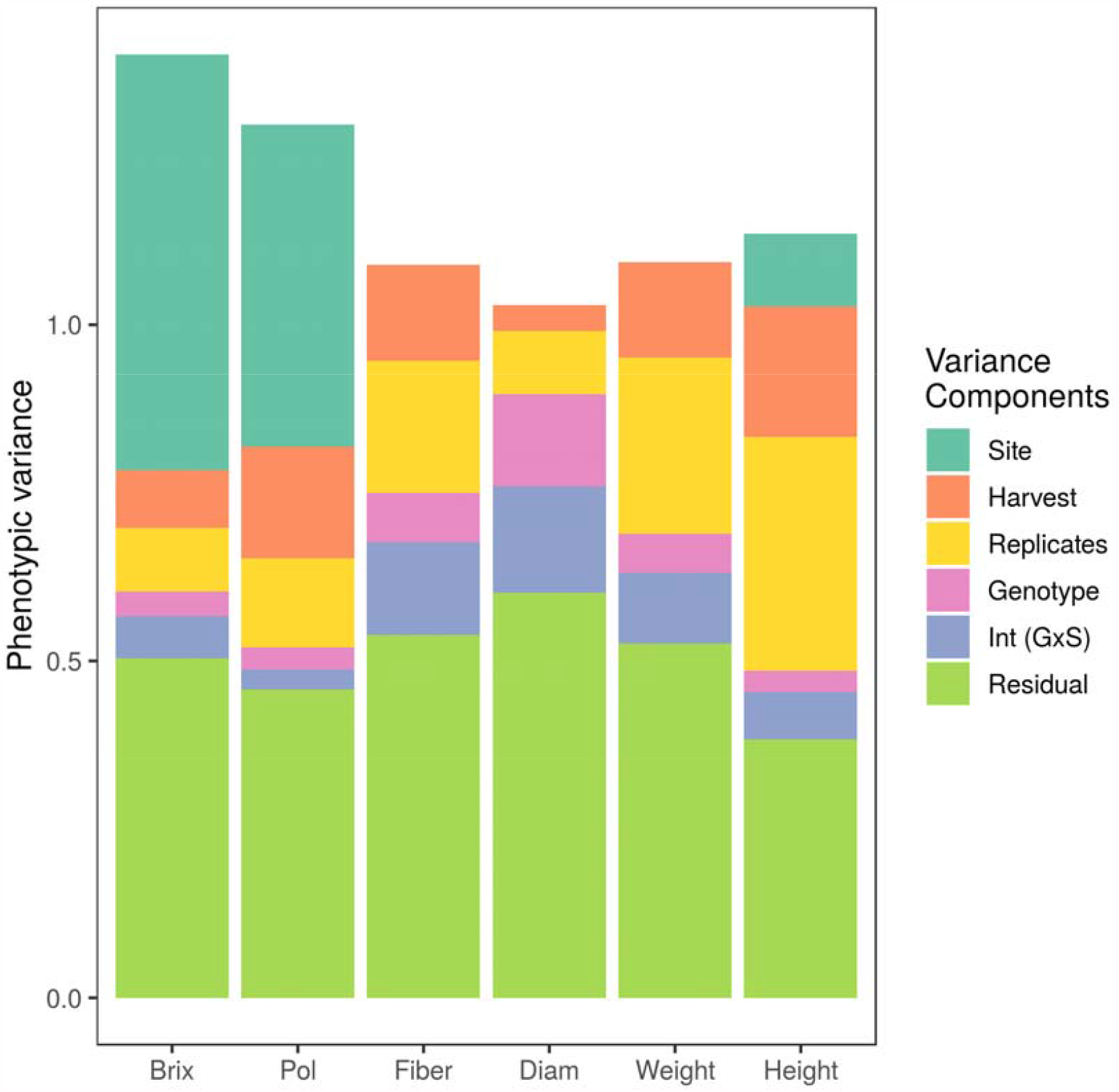
Phenotypic variance partitioning for soluble solids content (Brix), sucrose content (Pol), fiber percentage (Fiber), stalk diameter (Diam), stalk weight (Weight), and stalk height (Height). Variance components that are not shown had variance estimates very close to zero. Contributions of variances due to the effect of sites, harvests, replicates, genotypes, genotype × sites interaction (GxS), and residual variance are shown.

#### Sweet Potato

The predictive abilities of the genomic selection models in sweet potato were moderate to high and the distribution of predictive ability values were nearly equivalent between models and marker sets. Fig. 5 shows the distribution of the predictive ability values in the sweet potato dataset over different cross-validation runs of the G and G+D models when using all the makers with full ploidy and allele dosage information and using diploidized makers.

**Fig 5.**
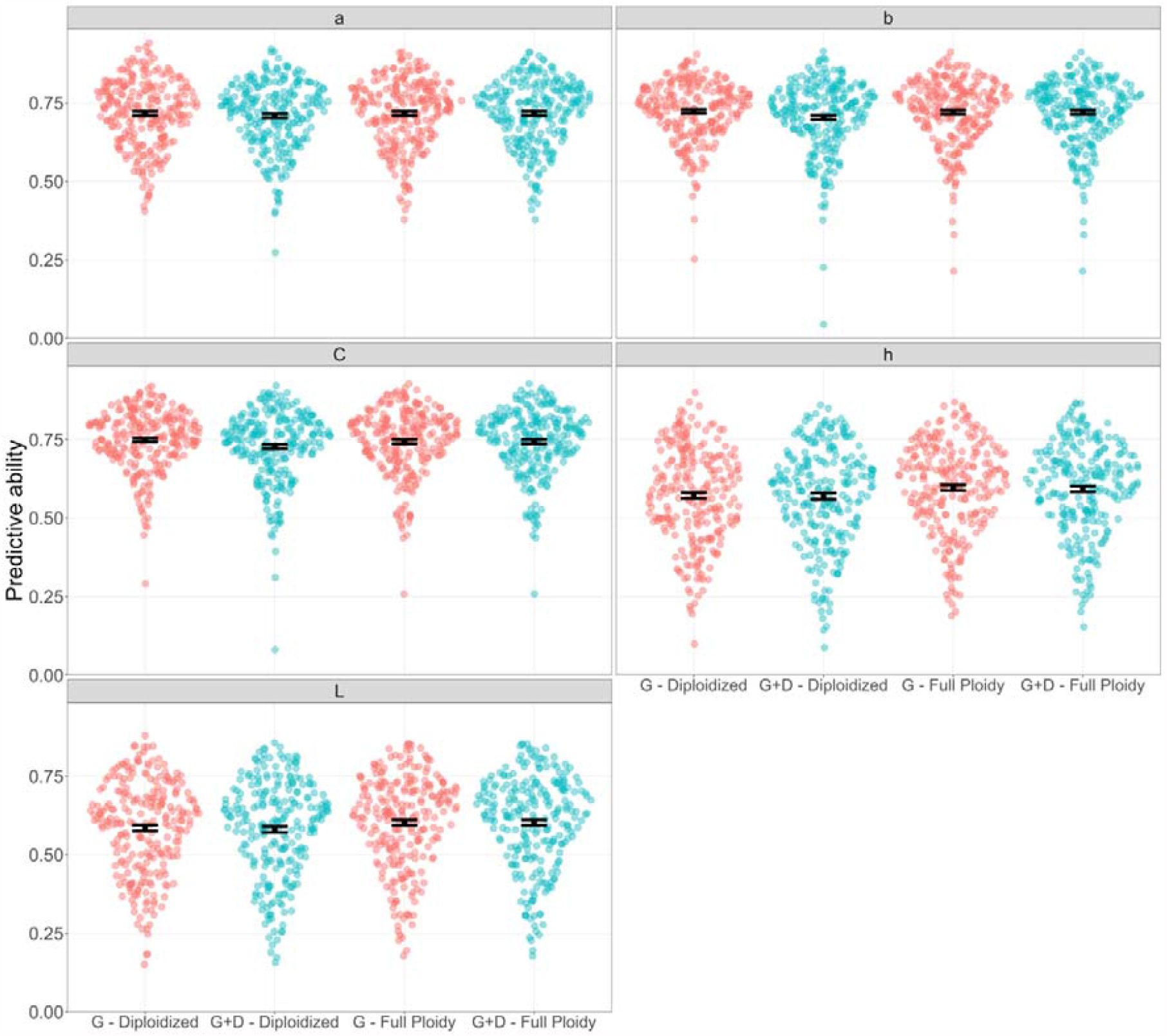
Distribution of the predictive ability values over different cross-validation runs of genomic selection in sweet potato. Values are shown when considering additive effects only (G) and considering additive and digenic dominance effects (G+D). Both models were compared when using markers with ploidy and allele dosage estimates (Full ploidy) and diploidized markers. The values are shown for stele colorimetry traits: green-red coordinate (**a**), the yellow-blue coordinate (**b**), color saturation (**C**), lightness (**L**), and hue angle (**h**). Mean and 95% confidence intervals are shown in black at the centre of each distribution.

The values of mean predictive ability for the green-red coordinate (**a**), the yellow-blue coordinate (**b**), and color saturation (**C**) were similarly high and barely differed between marker sets and models. The G model using diploidized markers, the G and G+D models using full dosage information had nearly equal mean predictive ability for all three traits: 0.72, 0.72, and 0.75 for **a, b**, and **C**, respectively. The G+D model using diploidized markers had slightly lower predictive ability values of approximately 0.71, 0.70, and 0.73 for a, b, and C, respectively.

For lightness (**L**) and hue angle (**h**) the mean predictive ability values were lower than for the other three traits. Predictive abilities were slightly higher when including the dosage information and did not differ whe dominance effects were icluded in the model. The G+D model using diploidized markers and markers with dosage information had nearly equal mean predictive abilites of aproximately 0.60 and 0.59 for **L** and **h**, respectively. The mean predictive abilites for the G model also did not differ between marker sets, with values of aproximately 0.58 and 0.57 for L and h, respectively.

### Simulations

In the simulated datasets the highest predictive abilities were achieved when including full ploidy and dosage information. Fig. 6 shows the distribution of the predictive ability values in the simulated datasets over different cross-validation runs of the G and G+D models when using all the makers with full ploidy and allele dosage information and using diploidized makers.

**Fig 6.**
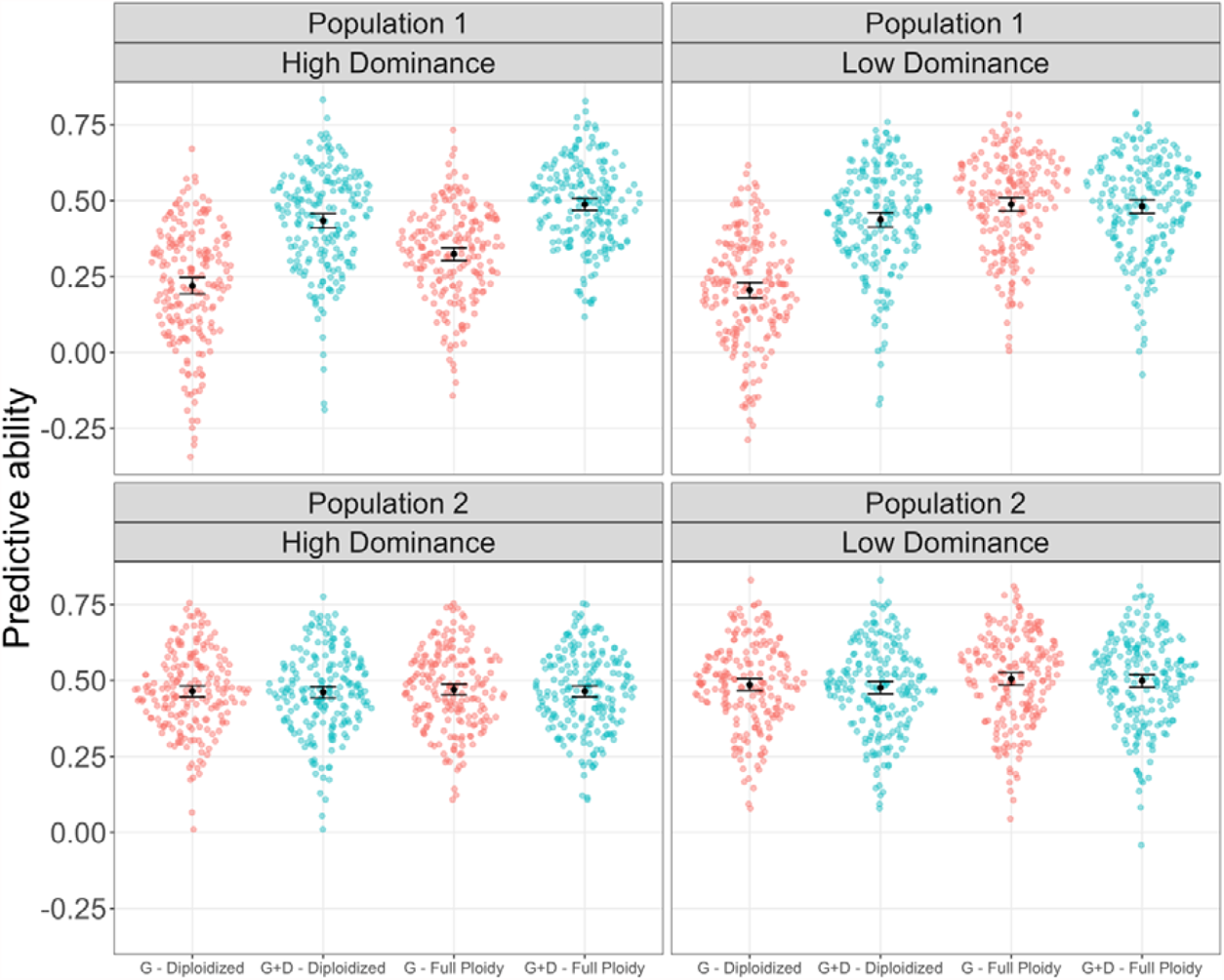
Distribution of the predictive ability values over different cross-validation runs of genomic selection in simulated datasets. Values are shown when considering additive effects only (G) and considering additive and digenic dominance effects (G+D). Both models are compared when using markers with ploidy and allele dosage estimates (Full ploidy) and diploidized markers. Simulated scenarios comprise populations with evenly distributed genotype frequencies (Population 1) and populations high number of homozygous and simplex markers (Population 2), either with low or high dominance. Mean and 95% confidence intervals are shown in black at the centre of each distribution.

When using dosage information, including digenic dominance effects was only advantageous under a high dominance degree and when the genotype frequencies in the population were more evenly distributed (Population 1). In this scenario, when using full ploidy markers, the G and G+D models had mean predictive abilities of 0.32 and 0.48, respectively. The mean predictive ability of the G+D model when using diploidized markers (0.43) was lower than that of the G+D model using dosage information. The G model using diploidized markers had the lowest mean predictive ability value (0.22).

For Population 1 with a lower dominance degree, when using full ploidy markers the mean predictive ability of the G+D model (0.48) was nearly equal but slightly lower than that of the G model (0.49). When using diploidized markers there was a clear advantage of including dominance in the models, with mean predictive abilities of 0.20 and 0.43 for the G and the G+D models, respectively.

When the frequency of heterozygous genotypes in the population was low (Population 2) the values of mean predictive ability for the different models and markers were similar in all simulated scenarios. For the low dominance degree level, the mean predictive abilities were approximately 0.50 for both the G and G+D models using dosage information, and 0.49 and 0.48 when using diploidized markers. For the high dominance degree level, the mean predictive abilities were approximately 0.47 for both the G and G+D models using full dosage information, and approximately 0.46 with the less informative diploidized markers.

## Discussion

We present our discussion in two sections. First, we discuss the results we obtained implementing genomic prediction in the sugarcane and sweet potato datasets. Second, we discuss the results we obtained with the simulated datasets and compare those with what we obtained with the real data. In both sections, we also show how models could potentially be improved to address the limitations in our study.

### Genomic prediction in sugarcane and sweet potato

The values of prediction ability for sugarcane were low, while for sweet potato we were able to obtain moderate to high values of predictive ability. Regardless of the prediction ability magnitude, for both species there was no significant improvement in predictions when including allele dosage information or dominance effects in the model. For sugarcane, we believe the low heritability and the size of the population were the main reasons why prediction models had a low performance. In both species, the high number of homozygous and single dosage markers are likely playing a role in the low sensitivity of the models to including dosage information and digenic dominance effects.

We were able to obtain high-quality genotypic data in sugarcane. We identified 6,550 SNPs with high mean read depths, high posterior probability of genotypes and ploidy estimates. Our SNP set exceeds in marker count many genetic studies in sugarcane (Bundock *et al*. 2009; Gouy *et al*. 2013; Costa *et al*. 2016; Yang *et al*. 2017; Gutierrez *et al*. 2018). However, the phenotypic variance partitioning analysis showed that, for all traits, most of the variation observed in the field experiments did not stem from differences between the individuals in the F_1_ progeny, as the variance components associated to the effect of genotypes and genotype × environment interactions had low magnitude in comparison to other experimental sources of variation. These low values of genotypic variability resulted in low to intermediate values of heritability, which in turn are usually associated with lower predictive ability (Combs and Bernardo 2013; Lian *et al*. 2014). For all of the traits we evaluated, several studies have reported higher heritability coefficients when analysing data from sugarcane cultivar trials (Milligan *et al*. 1990; Gravois and Milligan 1992; Tena *et al*. 2016). This indicates that implementing genomic selection in sugarcane is likely to be more advantageous than our results may suggest. Higher values of genomic predictive ability in sugarcane have been reported by Gouy *et al*. (2013), Deomano *et al*. (2020) and Hayes *et al*. (2021).

The small training population size in the sugarcane dataset might also be playing a key role in explaining the low values of predictive ability we observed. This idea is supported by comparing predictive abilities of the models when including or not including digenic dominance effects. For most of the traits there was a small reduction in the predictive ability when digenic dominance effects were included. Including digenic dominance effects results in estimating three additional parameters (Eq. 1), thus requiring more observations for accurate estimates to be obtained (Button *et al*. 2013). With a small population size, the estimates of dominance effects were likely not accurate, and the predictive ability of the model decreased.

In both datasets a large proportion of the SNP calls corresponded to either homozygous or single-dosage genotypes. In breeding populations, this can either occur when the polymorphisms genotyped represent relatively recent mutations in the genomes or when selective pressure has led to the near fixation of genotyped loci. In highly polyploid species such as sugarcane and sweet potato, even with very intense selective pressure, the fixation of favorable alleles is extremely difficult as deleterious alleles may have a high number of copies. Hence, the presence of recent mutations is the likely explanation for the genotype frequencies we observed.

This low frequency of higher-dosage genotypes is potentially masking the advantages of including allele dosage information in genomic selection models. As mostly only one class of heterozygous genotype is present, the marker sets with dosage information are not much more informative than their diploidized counterparts. We verified the effect of the low number of heterozygous classes in our prediction models by using simulated datasets, and we showed that it indeed affects the sensitivity of prediction models to the two different marker sets. In the following section we discuss these results more thoroughly.

### Genomic prediction in simulated datasets

The simulation results demonstrated that genomic prediction including allele dosage information and digenic dominance effect leads to higher predictive abilities only when there is a substantial presence of different heterozygous genotypic classes in the population (Population 1). As mentioned in the previous section, this is likely the main reason why predictions did not improve when including allele dosage information for the real datasets we used in this study. When the simulated populations had a higher frequency of homozygous and simplex genotypes (Population 2), and therefore a similar genotype distribution to the sugarcane and sweet potato datasets, we observed the performance of genomic prediction models to also be invariant to the inclusion of allele dosage and dominance effects.

With this, the results demonstrate that the simulated populations are a good proxy for better understanding the results we obtained in the real datasets. In addition to that, the highest value of mean predictive ability, obtained when including allele dosage information and digenic dominance effects, matched the value of the simulated broad- sense heritability of 0.5. Hence, the variance explained by the predicted additive and dominance effects fully captured the variance explained by the true genetic effects. This indicates that the model is capturing true genetic signals and is unlikely to overfit due to noise in the training data. This also highlights the low heritability values being the main culprit on the low predictive abilities observed in the sugarcane datasets.

Our results also demonstrate that the use of diploidized markers is a good alternative when allele dosage estimates are not available. This is true even with a sizeable presence of different heterozygous genotypic classes (Population 1). In these simulated scenarios, we observed that the performance of the G+D model using diploidized markers nearly matched the performance of the G+D model when considering allele dosage information, regardless of the dominance level. This is important because when using genotyping-by-sequencing techniques in autopolyploids, accurate genotype calls with allele dosage demand a high sequencing depth (Uitdewilligen *et al*. 2015; Bastien *et al*. 2018). When only low-depth sequencing data is available, making diploidized genotype calls can be an efficient way of using the data without having to obtain allele dosage estimates (Matias *et al*. 2019). Our results show that, in these situations, if dominance effects are included in the prediction model, the performance loss for using diploidized markers is not drastic.

Including dominance in the model is also important when using the allele dosage information. However, in this case, including digenic dominance effects is only advantageous when the dominance degree is high. When allele dosage information is included and the dominance degree is low, the G model performs just as well as the G+D model. In contrast, under high dominance degree levels, the performance loss when using the G model rather than the G+D model is significant. To date, little is known about the magnitude of the dominance gene action that is present in the traits of highly autopolyploid species. More research is still needed for breeders to have an estimate of the dominance level of traits in autopolyploid breeding populations. In the current context, our results show that the G+D model should be preferred, as it is the best performing model regardless of the dominance level.

Generally, autopolyploid crop varieties are clonally propagated and the genotypes in vegetatively propagated crops are typically heterozygous (Grüneberg *et al*. 2009). The genetic value of heterozygous genotypes is a function of additive and non- additive gene action (Falconer and Mackay 1996). Non-additive gene action comprises both dominance and epistatic effects. For clonally propagated species, both additive and non-additive gene action are transmitted across generations in the selection process (Bernardo 2010). Therefore, genomic selection models for cultivar selection in these species should aim to include both dominance and epistatic effects. The importance of including dominance effects in genomic models for clonally propagated crops has also been demonstrated for selection of parents in recurrent selection breeding programs (Werner *et al*. 2020). In this study, we investigated only two of many possible ways of including dominance effects in prediction models for highly polyploid species.

Is important to notice that we validated out models using simulated phenotypes consisting of only additive and digenic dominance effects. We were able to demonstrate that our fully informative dosage-aware analysis performs better than other simpler genomic prediction models when it comes to these two simulated effects. However, higher order dominance effects (i.e., interactions between more than two alleles) could also be present in autopolyploid species; hence further improvements in predictions could be achieved by expanding genomic prediction models to include these effects. Moreover. it is still unclear how much of the genotypic variation in highly autopolyploid species is explained by digenic dominance effects. In autotetraploids, Endelman *et al*. (2018) and Amadeu *et al*. (2020) have observed digenic dominance effects to explain only a small portion of the genotypic variance. In their case, there was little advantage to including digenic dominance effects in genomic predictions.

## Conclusion

We showed that estimates of ploidy and allele dosage can improve genomic selection in highly polyploid species. This is mostly true when there is a substantial number of heterozygous genotypes in the population. When the frequency of heterozygous genotypes in the population is low, such as in the sugarcane and sweet potato datasets, there is little advantage in including allele dosage information in the models. Our simulation results also show that using diploidized markers in the absence of allele dosage estimates can nearly match the performance of fully informative marker sets. However, this is true only when including dominance effects in the genomic prediction models. With the full dosage information available, digenic dominance effects can significantly improve genomic prediction, provided that the trait being predicted has a high mean dominance degree and that the population has a high frequency of heterozygous genotypes.

## Declarations

### Availability of data and code

The sugarcane and sweet potato datasets as well as the code for obtaining genomic covariance matrices of additive and digenic dominance effects can be found on the github repository https://github.com/Lorenagb/GS_HighlyPolyploid. The code for generating all four simulated datasets can also be found on the same github repository.

## Authors Contributions

LGB, APS and GRM conceived the study. APS provided the genotyping by sequencing raw read data for the sugarcane population. LGB and VHM performed the SNP calling in the sugarcane and sweet potato datasets. LGB expanded and implemented the genomic selection models and designed the plant breeding program simulations. All authors read and approved the manuscript.

