## Supplementary Material 1 for "Genomic prediction with allele dosage information in highly polyploid species"

**Phenotypic analysis**

For phenotypic variance partitioning and to obtain estimates of heritability for the traits, the phenotypic values were first standardized and the following random linear model was used for each trait:

$$y_{ijkl}= \mu+g_{i}+s_{j}+ h_{k}+b_{l(jk)}+{(gs)}_{ij}+{(gh)}_{ik}+{(gsh)}_{ijk} +e_{ijkl},$$

where $\mu$ is the intercept, $g_{i}$ is the effect of genotypes, $s_{j}$ is the effect of sites, $h_{k}$ is the effect of harvest, $b_{l(jk)}$ is the effect of replicates within sites and harvests, ${(gs)}_{ij}$ is the effect of the genotype × site interaction, ${(gh)}_{ik}$ is the effect of the genotype × harvest interaction, ${(gsh)}_{ijk}$ is the effect of the genotype × site × harvest interaction, and $e_{ijkl}$is the residual effect.

The estimates of heritability were obtained using:

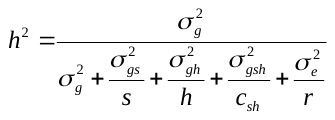
,

where
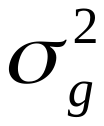
,
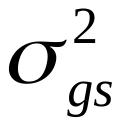
,
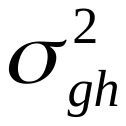
,
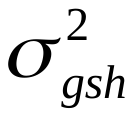
, and
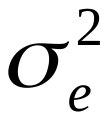
are the genotypic variance, the variance of the genotype × site interaction, the variance of the genotype × harvest interaction, the variance of the genotype × site × harvest interaction, and the residual variance, respectively. The values
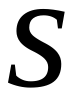
,
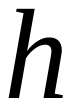
,
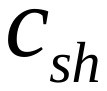
, and
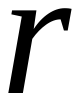
correspond to the number of sites, the number of harvests, the number of combinations of sites and harvests, and the total (combined) number of replicates of both experiments, respectively.

**Simulated datasets**

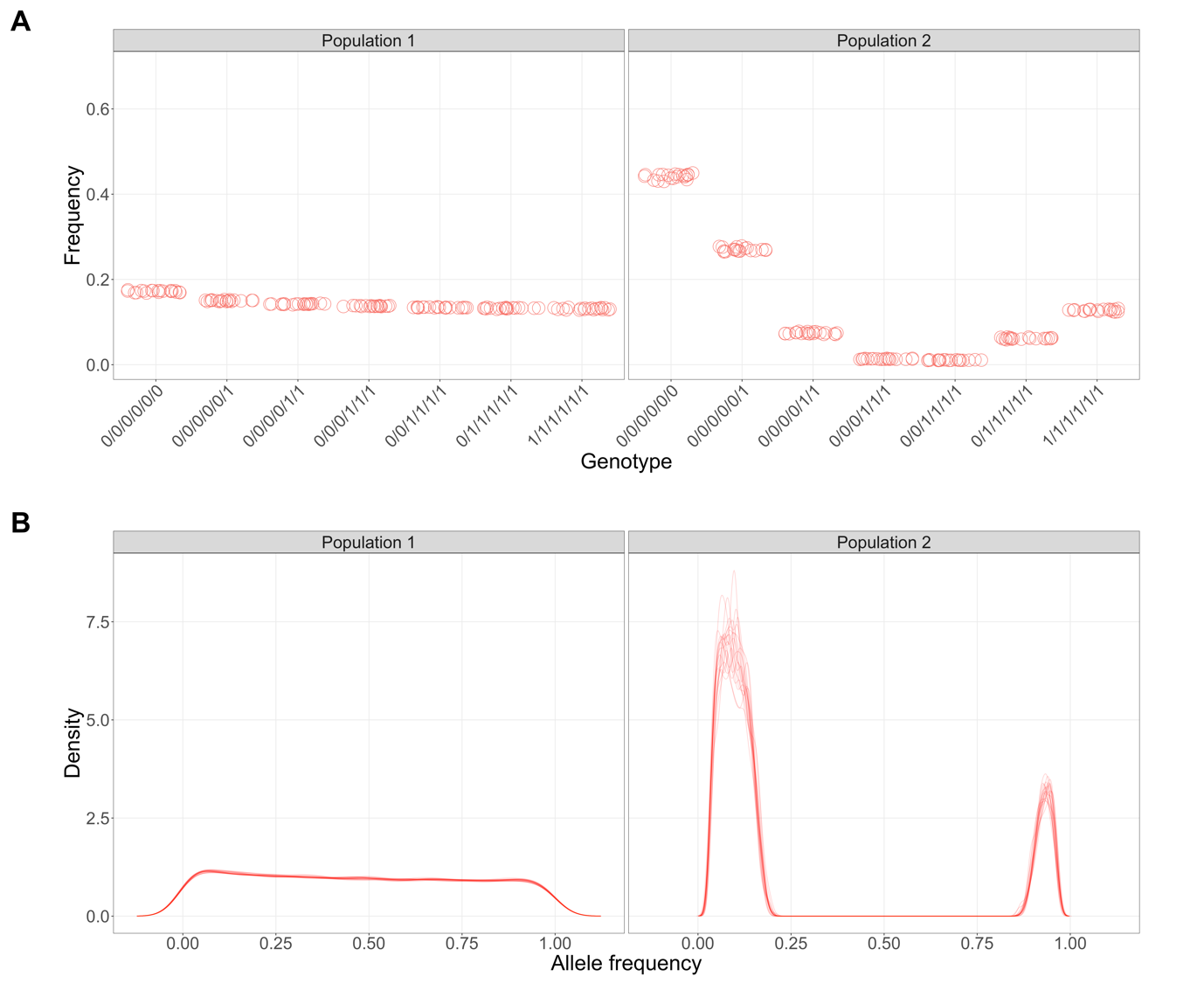

Figure S1.1. Distribution of SNP genotype (A) and allele (B) frequencies in simulated populations with a low mean dominance degree (0.3). Populations had either genotype frequencies more evenly distributed (Population 1) or a higher frequency of homozygous and simplex genotypes (Population 2). Different circles and lines correspond to different replicates of the simulations

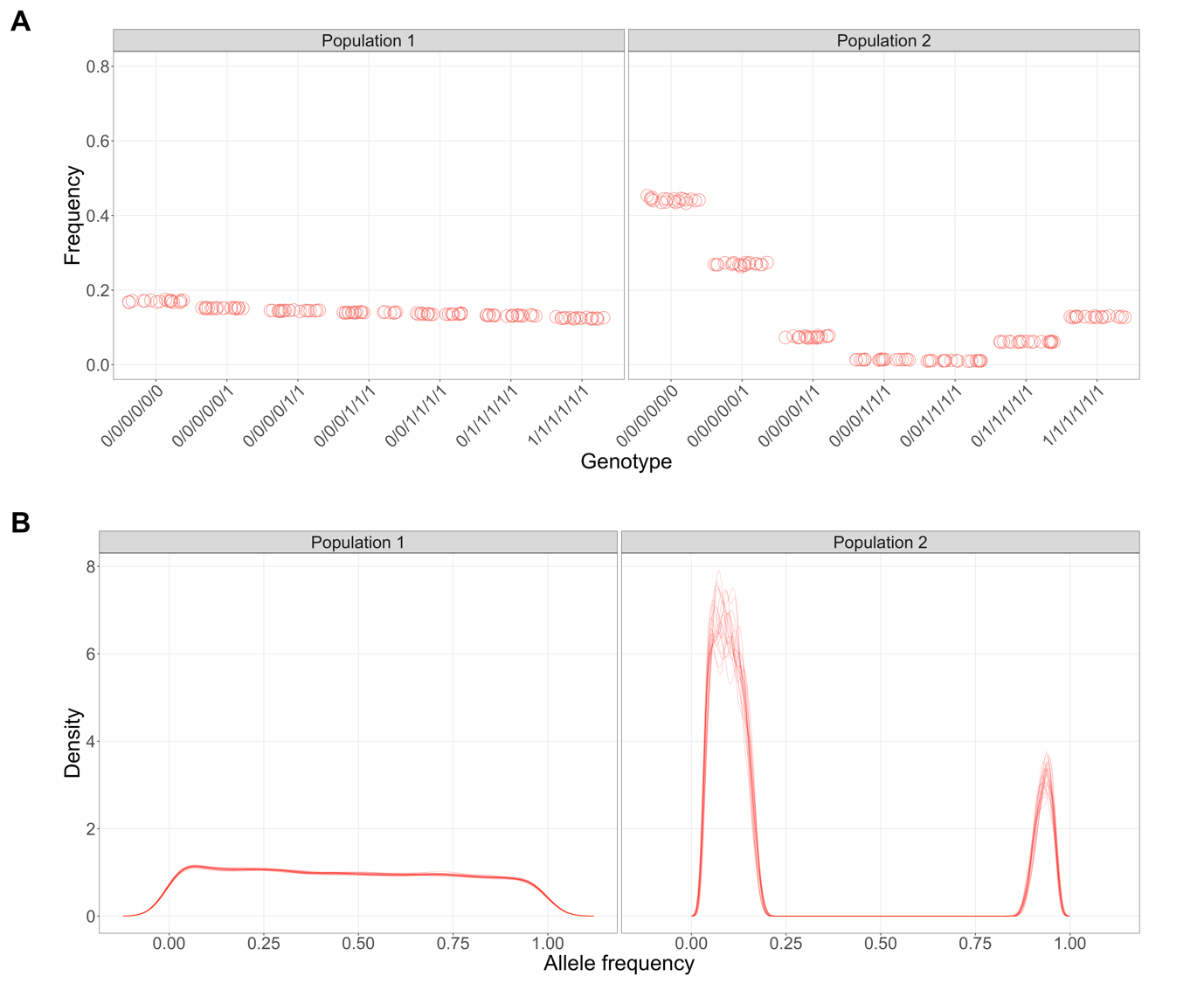

Figure S1.2. Distribution of SNP genotype (A) and allele (B) frequencies in simulated populations with a high mean dominance degree (1.0). Populations had either genotype frequencies more evenly distributed (Population 1) or a higher frequency of homozygous and simplex genotypes (Population 2). Different circles and lines correspond to different replicates of the simulations
